# Sponge Physioecology On Moorea, French Polynesia: Local Distribution And Filtration Efficiency Of *Lamellodysidea* Under Varying Temperatures

**DOI:** 10.1101/2020.01.14.906990

**Authors:** Lily Leveque-Eichhorn

## Abstract

This physio-ecological study investigated the physiology and distribution of *Lamellodysidea sp.* in Moorea, French Polynesia. Specifically, its distribution was described across three reef types—fringing, mid-barrier, and barrier reefs—as well as across sites between Cook’s Bay and Opunohu Bay. Additionally, filtration experiments were conducted to test how temperature impacts filtration efficiency. This is informative when predicting how future ocean temperatures are going to affect sponge’s success and distribution. Sponge abundance was found to decrease from the fringing reef out to the barrier reef, with the highest number of sponges in the fringing reef, less in the mid-barrier reef, and none found in the barrier reef. Sponges were also unevenly distributed across sites, with sponge abundance clearly increasing as you move away from Cook’s Bay. Together, these data show that sponges demonstrate habitat preference that is related to their physiological tolerances. Filtration data showed that over a 3-hour period sponges increase their physiological output when introduced to environments 3-6 degrees Celsius above normal. This suggests that as ocean temperatures warm, sponges are likely to increase their filtration efficiency, thus increasing their ecological role as filter feeders, or struggle to survive at this new level of physiological function.

## Introduction

Coral reefs are among the most diverse ecosystems in the world and the most diverse marine ecosystem per unit (Bellwood and Hughes 2005, Knowlton et al. 2010). Reef communities are made up of complex species interactions with some species contributing disproportionately to overall reef health as ecosystem service providers. Naturally, reefs experience disturbances on a regular basis in the form of large storms, fluctuating temperatures and changes in other abiotic conditions. In face of anthropogenic pressures such as climate change, overfishing, and decreasing water quality, disturbance events are occurring more frequently and severely (Feary et al. 2007). This raises questions about how ecologically important species will cope with these increasing stressors and what that means for overall reef function.

Sponges contribute to the functioning of coral reefs in unique and significant ways by participating in spatial competition and exclusion with corals and other species (Rutzler 1970, Thacker et al. 1998), aiding reef cementation (Wulff 1984), and influencing nutrient cycling within the reef (Bayer 2008, Yahel 2003, Reincke and Barthel 1997). As rigorous filter feeders, sponges play an essential role in trophic linkages as they regulate production and availability of dissolved nutrients, which in turn impacts primary productivity and thus the rest of the food chain (Maldonado et al. 2012). Many animals, including spongivorous fishes and the hawksbill sea turtle, depend on sponges as a food source, further demonstrating their importance in reef ecosystems (Leon and Bjorndal 2002). Authors have proposed that sponges may become the dominant animal in reefs in coming years as they are less sensitive to ocean acidification compared to corals (Bell 2013). For these reasons it is essential to have a baseline understanding of sponge’s current distribution and abundance, as well as work to deepen our understanding of their physiology in light of environmental changes.

As the first known metazoans, evolving 760 mya, sponges reflect the biology of the first forms of multicellular life on earth (Brain 2012). Sponges are made up of special cells called choanocytes, which beat their flagella in order to sweep water through the sponge where dissolved organic matter (DOM) and particulate organic matter (POM) is digested via phagocytosis. All sponges are thought to reproduce both sexually and asexually and most sponges are monecious, meaning they contain both sexes.

Despite their ecological and evolutionary importance, large gaps of knowledge still exist regarding sponge’s distribution, basic taxonomy, and physiology (Wörheide and Erpenbeck 2007, Thomassen and Riisgard 1995). As our planet undergoes rapid environmental change—by some estimates, the sixth mass extinction (Wake 2008)—it is critical to understand current species’ distribution and abundance so that we can align our conservation efforts in order to preserve biodiversity hotspots and endemism as well as track population shifts through time.

In Moorea, French Polynesia, sponges have been documented on the fringing reef, mid-barrier area, and barrier reef (Moorea Biocode Project Nov. 11^th^ 2019, Desmet 2009, Murphy and Wright 1998). Yet, detailed records of sponges’ distribution, abundance, and population characteristics in respect to these three reefs remains largely unknown. My study focuses on describing the distribution and physiology of one especially valuable sponge genus—*Lamellodysidea.* Understanding *Lamellodysidea* sponges’ distribution between Cook’s Bay and Opunohu Bay in relation to fringing, mid-barrier, and barrier zonation is important when considering the habitat preferences and physio-ecological function and tolerances of this group of sponges. As changes in abundance and distribution of *Lamellodysidea sp*. would likely be coupled with large-scale ecological changes due to their robust ecological role, it is important to establish a record of these sponges so that future studies may be able to look at reef shifts over time.

Furthermore, most of our existing knowledge about sponge physiology is based on information from cold-water sponges, which may function very differently compared to tropical sponges (Riisgard et al. 1993). For this reason, I incorporated a physiological experiment into my study, which tested how temperature impacts *Lamellodysidea sp.* filtration efficiency. The goal of this part of my project was three-fold: (1) To understand how sponges might deal with warming water temperatures due to climate change (2) To see if their physiological tolerances in relation to temperature can be predictive in terms of sponge’s distribution given that water temperature gradients exist within the lagoon (3) To look at their ecological role at different temperatures. My research had two parts: (1) A field survey to investigate *Lamellodysidea sp.* distribution and abundance relative to reef type (fringing, mid-barrier, and barrier) and site (2) A filtration experiment, to understand how temperature impacts sponge filtration efficiency, which was treated as a proxy for physiological function. Given that that barrier reefs experience more wave action and generally have harsher conditions, I expected fringing and mid-barrier reefs to host denser sponge populations compared to the barrier habitats. Additionally, I expected sponges to exhibit shallow depth preference, given that many sponges contain photosymbiotic communities and thus need light. For my lab experiments, I hypothesized that, similar to cold-water sponges, *Lamellodysidea sp.* would show increased filtration efficiency at higher temperatures compared to sponges at ambient temperature (Riisgard et al. 1993).

## Methods

### Study site

This study was conducted on the island of French Polynesia from October 12^th^ to November 22^th^ 2019. Four sites were chosen between Cook’s Bay and Opunohu Bay on the north side of the island. Sites were spaced relatively evenly between these two bays and were selected based on access. Preliminary observations were made to categorize fringing, mid-barrier, and barrier habitats, each of which contained three transects. A stratified random design was employed to lay out 36 50 m transects total, 9 at each site at a total of 4 sites.

Thus, each habitat was represented by 12 transects and each site by 9. I used basic traits for habitat characterization which, along with approximate distance from shore, allowed me to identify these reef zones. See Appendix for detailed habitat descriptions.

### Field study

Surveys were conducted to characterize the distribution of *Lamellodysidea sp.*, and to test whether distribution differed between the three habitat types (fringing, mid-barrier, and barrier reefs). For each transect, latitude and longitude were recorded using the Altimeter GPS App. Additionally, five water depth measurements were taken evenly along each transect and averaged for the overall water depth of the transect. I also made detailed observations of habitat type at each transect. When I encountered a sponge along a transect, I recorded the following: sponge size, color, morphology (encrusting, encrusting/tall, tall), location, the depth from the sponge to the water surface, and what type of substrate the sponge was attached to. It was also noted whether or not any sponges were clumped, which was defined as sponges growing less than 0.3 m away from each other.

All analyses were conducted in R (R Core Team 2019) as implemented in RStudio (RStudio Team 2018) and alpha was set to 0.05. I used a Kruskal-Wallis test to look at whether sponge abundance varied by habitat (fringing, mid-barrier, barrier). In this case habitat is being treated as an explanatory variable and sponge abundance is a response variable. A Kruskal-Wallis test was also used to look at how size (response variable) varied by habitat (explanatory variable). An analysis of variance (ANOVA) was employed to test the significance of study sites on sponge abundance. A linear regression model was used to test how sponge abundance (response variable) varied with sponge depth (explanatory). I also employed a mixed effects model in the R package and lme4 to understand the influence of habitat on sponge depth (fixed effects) with the random effect of transect (Bates et al. 2015).

### Filtration experiment

Experiments were conducted at the Gump Research Station between November 11^th^ and 20^th^. The goal of these experiments was to measure *Lamellodysidea sp*. ability to filter water at varying water temperature regimes. This is a basic way to look at physiological stress of a filter feeder at different temperatures. Animals for all replicates were collected in the same general area where preliminary observations showed sponges to be large and abundant. See Appendix C for GPS point. For each trial, sponges were collected from the same sponge aggregation in order to eliminate individual filtration variance as a confounding factor. Three sponges were required for each trial so either 3 or 6 sponges were collected per trip for a total of 18 sponges for all 6 replications. After sponges were collected, they were held in a non-shaded water table between 4 to 24 hours before experiments were conducted. To ensure that sponges across treatments were the same size, their volume was measured (as water displaced from a 1000m beaker) and adjusted prior to each experiment. Since sponges are known to be very sensitive to air exposure, sponges were only taken out of the water for a few seconds to measure their volume (Desmet 2009). Four tanks were set up, one for each treatment—no sponge control, temperature control, 32 degrees and 35 degrees. The purpose of the no sponge control treatment was to account for small variances due to light changes throughout the experiment and to have a baseline measurement of the accuracy of my measurement design. The temperature control treatment was useful to look at filtration under normal ambient water temperatures. 32C is approximately 3 degrees above ambient and is useful when thinking about future water temperatures. 35 C was chosen to attempt to test the upper physiological limit of *Lamellodysidea sp.*

Sponges were placed in 3 tanks with 5ml of powdered milk and equal amounts of water, which had been heated using a Deltatherm Interpet aquarium heating system to appropriate treatment temperature. The powdered milk “clouded” the water, making turbidity change detectable as sponges non-selectively filtered the water. Experiments took place over a three-hour period during which turbidity was measured at regular 15 and then 30-minute intervals. A secchi disk was mounted to the bottom of a graduated cylinder and was used to measure turbidity. All measurements were taken in the shade in order to minimize light variance between samples.

For filtration analysis, three full and three reduced generalized linear mixed effect models were built to look at the effect of multiple fixed factors (time since start, treatment, and interaction between time since start and treatment) on water turbidity. These mixed linear models were performed with the R packages lmerTest and lme4 (Kuznetsova et al. 2017, Bates et al. 2015). Sponge number, which was distinct between trials, was treated as a random effect. ANOVA tests were then used to compare full and reduced models and to parse out which fixed explanatory factors most influenced turbidity change—response factor.

## Results

### Field study

Sponges were observed in both fringing reefs and mid-barrier reefs across all sites, yet not found in any barrier sites and thus not represented in fig 2 and 3. Sponge abundance was higher in fringing reefs compared to mid-barrier reefs. Overall, abundance varied by reef type significantly (p = 0.005495 Kruskal-Wallis test, chi square = 10.408, df = 2). Sponge abundance also varied between sites with marginal significance, increasing from site 1 through 4 as shown in figure 3 (p-value = 0.044 at sign = 0.01 ANOVA).

**FIG. 1.**
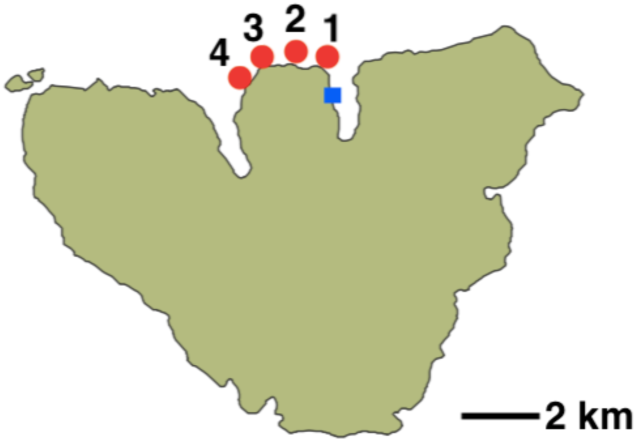
Map of Moorea, French Polynesia, showing four study sites (red dots), with each site containing a fringing, mid-barrier, and barrier habitat, with a total of 9 transects per site. The blue square notes the Gump Research Station in Cook’s Bay.

**FIG. 2.**
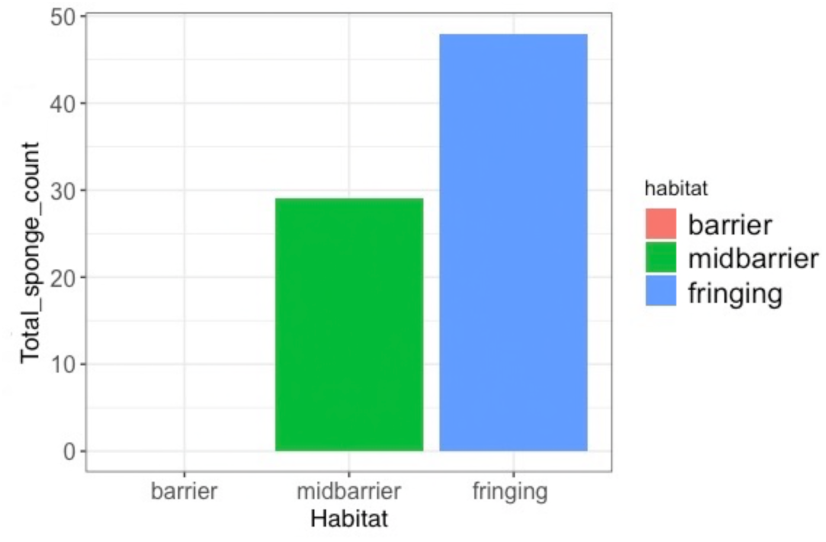
Describes the relationship of sponge abundance with reef type. Total sponge count accounts for all 77 sponges seen across all 36 transects.

**FIG. 3.**
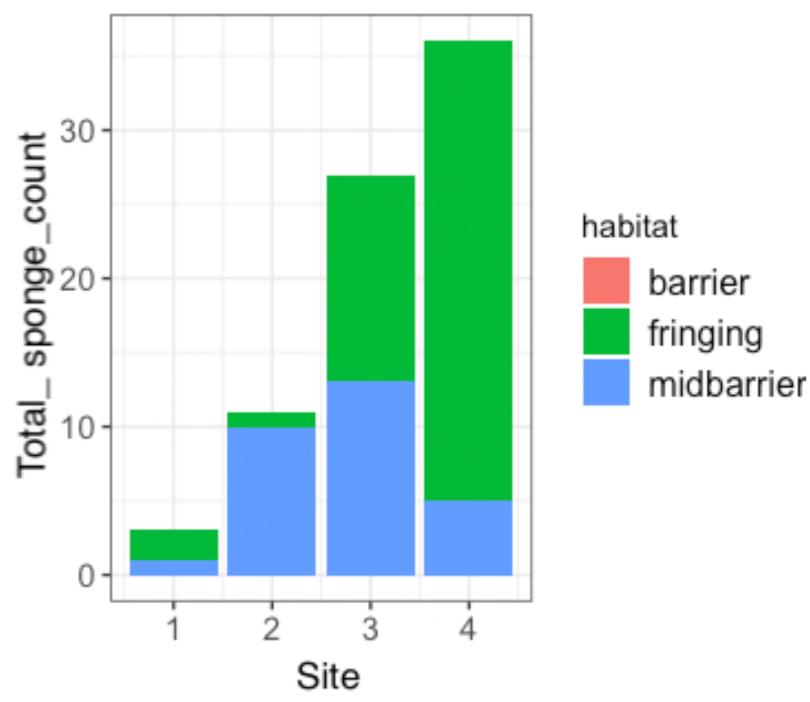
Describes total sponge abundance relative to site. In total, 77 sponges from the 36 transects are represented here.

Sponge abundance and depth (m) did not seem to be correlated, although Fig 4 shows two general depth-peaks where sponges were most abundant. These peaks likely correlate with common water depths of these two habitats. ANOVA analysis showed strong correlation between depth and habitat, further supporting this (p> 0.0001). Thus, sponge count seems to vary more with habitat type than with depth. Fig 5. shows a slight difference in average sponge size, measured as cm^2^, however both an ANOVA and Kruskal test determined that this difference is not significant (p>0.05).

**FIG. 4.**
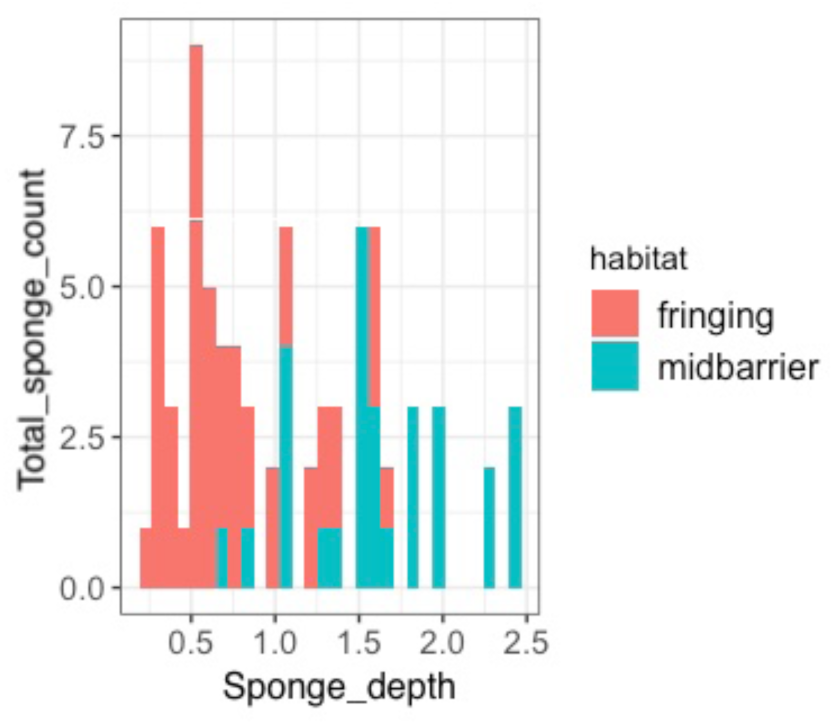
Sponge abundance as measured by total sponge count in relation to sponge depth (m) across all 36 transects. In total, 77 sponges are represented here. Two peaks can be observed. One at approximately, 0.5 and the other at 1.5 m.

**FIG. 5.**
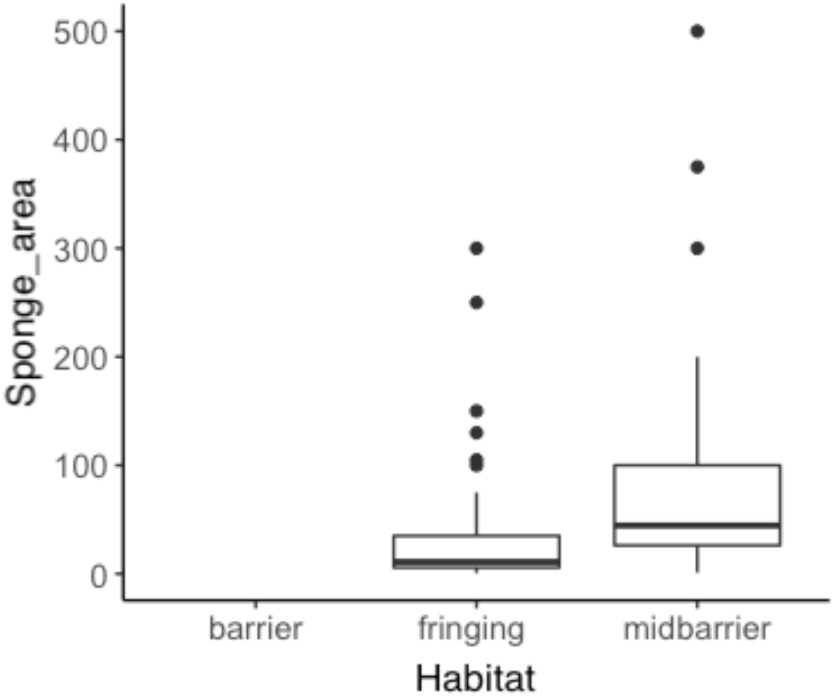
This figure depicts the relationship between sponge size, measured by sponge area in cm2, and habitat type. In total, 47 individual sponges are in the fringing reef and 26 are in the mid-barrier reef. Note that 3 outliers in the mid-barrier reef were omitted from this figure.

Note that, for the figure, three outliers were omitted; however, no data points were excluded in the statistical analysis. Clumped versus non-clumped sponge aggregate form was measured for each sponge and 45.5% of all sponges were found to be clumped. Additionally, 59.7% of sponges were found to be white/grey, and 40.3% white/yellow. Furthermore, encrusting growth form was dominant with close to 80% of observed sponges exhibiting this morphology. See appendix for details on the characteristics.

### Filtration experiment

ANOVA tests were conducted to determine the significance of multiple variables in relation to clearance rate, measured as turbidity change over time. There is a discernable difference in clearance rate between the two warm experiments, 32C and 36C, and between the control treatments. Indeed, an interaction variable testing the interaction between time and treatment carried statistical power (p < 0.001). This shows that, over time, there is a higher filtration rate at higher temperatures. However, overall treatment effect did not have statistical power (p> 0.05). I was not able to reject my null hypothesis which stated that overall turbidity would not change over time (p > 0.05). Fig. 6 shows differences between treatments and an overall increase in filtration over time for the two warm treatments. Note here that the units for filtration rate are volumes (ml) at which the secchi disk was no longer visible inside a graduated cylinder— meaning larger values correspond to higher clarity and lower turbidity. Mean turbidity is measured in ml. where values correspond to the volume at which the secchi disk could no longer be distinguished.

**FIG. 6.**
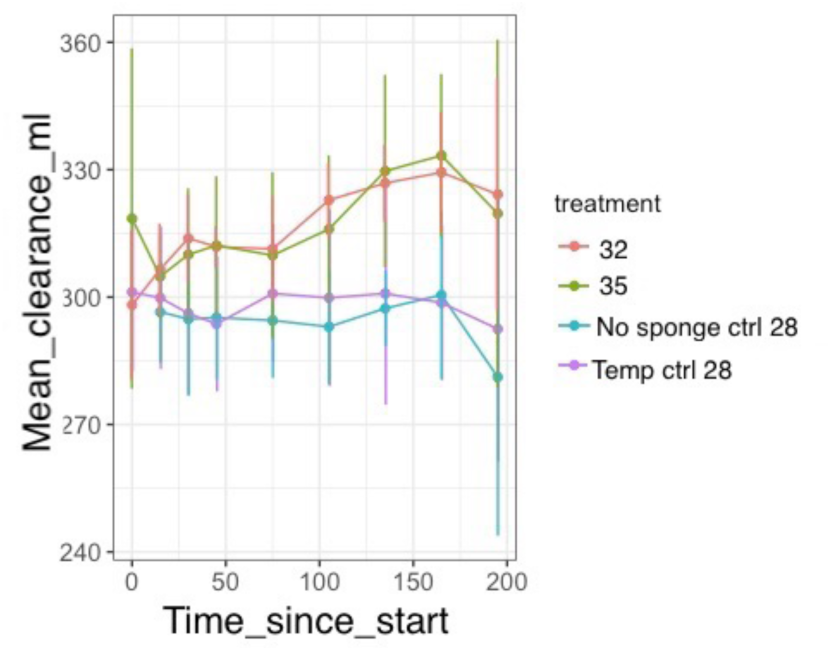
Water turbidity over time for four temperature treatments. Six experiment replications are compiled to form this graph

## Discussion

Sponges were observed to have an overall patchy distribution with patch sizes varying from approximately 25 m2 to over 100 m2. In these areas, sponges were often growing abundantly on dead coral heads, coral rubble, or directly on the sand. These groupings hosted sponges of varying colors and morphologies, which could suggest that multiple species of the genus *Lamellodysidea* are represented in each patch. Sponges, which were categorized as “white/yellow” and “white/grey” in this study, have been treated as different species in past studies. Desmet (2009) arbitrarily named sponges species A through C, characterizing them based on their morphology, color, texture, etc. However, without DNA analyses of these sponges, it is difficult to determine whether they are, in fact, different species or whether their color differences are simply due to varying symbiotic communities. My observations suggest the latter as individual sponges were observed to exhibit multiple morphologies and colors within one sponge aggregate. I propose that *Lamellodysidea herbacea* is likely the species of sponge I worked with, as backed by my own observations and BioPoly Val project’s specie’s description that noted color variation within *L. herbacea* (BioPoly Val Project Nov. 3^rd^ 2019). For the purposes of this paper, however, I referred to the group only at the genus level as without DNA analysis it is difficult to confirm identification to such a specific level. See Appendix for more details on identification and color variation.

Due to their sporadic distribution, it would likely be more efficient for future studies to select their study sites based on where sponges are found instead of randomly or otherwise. This would likely add substantial statistical power to one’s data as the volume of data would be much greater and help identify more sponge habitat preferences.

*Lamellodysidea sp.* is distributed differentially within and across reef types, which indicates habitat preferences and perhaps even physiological limitations of this sponge. Sponges were most abundant in fringing reef habitats, then mid-barrier reefs, and no sponges were ever found in the barrier reef. There are several potential explanations for this observed trend such as sponge temperature preference, wave action sensitivity, and substrate availability. Fringing reefs are much shallower compared to mid-barrier and barrier reefs and thus, generally, experience warmer and more fluctuating temperatures which organisms in these environments must cope with (Jimenez et al. 2008). As studies, including this one, have shown temperature to impact sponge physiology—as measured by filtration, reaggregation, etc.—it would not be surprising if sponges settled preferentially in habitats with a specific temperature range and are thus distributed in respect to a temperature gradient (Runzel 2016, Riisgard et al. 1993, Webster 2008). Substrate availability is another factor that may impact sponge distribution. Sponges were found attached most commonly to dead coral heads, which were observed to have higher macro-algae percent cover closer to the algal ridge. This may discourage sponge recruitment in the barrier reefs as less substrate is available and free of macro-algae. Wave action and nutrient availability are two other factors potentially leading to the differential sponge distribution across reef types. Sponges have been shown to be affected by wave action as they modify their morphology, increasing spicule count and size to favor a sturdier form, in areas with higher wave action (Palumbi 1986). Additionally, the distribution of *Halichondria panicea*, a temperate demosponge, has been observed relative to a wave action gradient, where sponges were not found in areas of high wave action (Palumbi 1986). Similarly, *Lamellodysidea sp.* may be affected by wave action and thus not settle in areas where this poses extra physiological challenges.

Reproductive strategies can also have a profound impact on species distribution. Population genetics work on *Lamellodysidea* would elucidate to what degree sponge reproduction and larvae settlement determine sponge distribution. Field observations have shown that after sponge-larvae is released from its mother-sponge, it can reach “competence”, defined as the necessary physiological and morphological state larvae need to exhibit in order to settle, just minutes after being released (Lindquist et al. 1997). The observed patchy distribution of *Lamellodysidea* may then be less about its habitat preferences and more about a quick dispersal time. Based on their distribution, it would not be unreasonable to hypothesize that *Lamellodysidea* larvae reach competence quickly and thus settle close to mother-aggregates. However, further investigation into sponge reproduction and dispersal is necessary in order to make any solid assertions about the role of these processes on observed distribution of *Lamellodysidea sp.* It is worth noting that asexual reproduction could also be partly responsible for these groupings as small pieces of sponge bud off and form new colonies. Roughly half of all sponges observed in my transects were clumped, which may indicate high levels of asexual reproduction within these larger patches. Genetics work would be needed to confirm this. Furthermore, genetic studies could help us understand which reproductive strategy is dominate and how this reproductive method may be partly responsible for the observed distribution.

Differences in sponge abundance were also found between sites, with site 1 having the lowest abundance and increasing to site 4. Cook’s Bay is much more commercially impacted compared to Oponohu, with more housing developments and agricultural activity. In 2006 a land use assessment of Moorea reported that soil pollutant levels around Cook’s Bay were between “moderate” to “very high” whereas soils around Opunohu were “minimally” to “somewhat” impacted (Duane 2006). With sediment runoff four times higher in the Poa Poa watershed compared to the Opunohu watershed, it is obvious that a water quality gradient exists between Cook’s Bay and Opunohu, with water quality improving as you move away from Cook’s Bay (Duane 2006). The observed difference in sponge abundance between these two bays may be, in part, a result of sponge’s inability to succeed in areas with poor water quality.

Many sponges have been documented as having specific depth preferences (Pang 1973, Desmet 2009). *Lamellodysidea* sponges do not appear to display this as they were found in shallow waters not even 1m deep as well as water greater than 2.5 m. However, sponges were most commonly found at two depths, roughly 0.75 and 1.5, which correspond closely to the average water depth at fringing and mid-barrier reef types. Since “preferred” depth corresponds so closely with common depth at these reef types, it is likely that sponge abundance varies with habitat type more than with depth. However, depth preference could be one of the factors attributing to reef preference, especially since depth limits light and many species of sponges, including species in the genus *Lamellodysidea*, contain photosynthetic symbionts which require base levels of light penetration (Usher 2008, Freeman et al. 2016).

Understanding the physiology of *Lamellodysidea sp*. in respect to temperature variance can also be helpful when thinking about sponge distribution patterns relative to water temperature gradients in the lagoon. Experimentation showed that at warmer temperatures (32 and 35C) sponges were able to filter more efficiently over a three-hour period. This seems to indicate that temperature does, in some way, impact *Lamellodysidea* sponge’s physiology. Cold water sponges have been shown to increase their filtration rate as a physiological response to higher temperatures (Jorgensen 1949). My experiments suggest that *Lamellodysidea*, and perhaps warm-water sponges in general, also have this response to warmer temperatures. It would make sense then for sponges to live in an optimal temperature range, which would be reflected in their distribution.

Given the observed distribution of *Lamellodysidea*, it is worth considering the ecological implications of this distribution across reef types and sites. Sponges, as we’ve established, play a robust ecological role. They not only create broad trophic linkages by controlling nutrient load, but they are also a critical food source for spongivores (Bayer 2008, Leon and Bjorndal 2002). In areas where sponges are more abundant, like the fringing reef and Opunohu bay, nutrients levels are likely affected by sponges and thus primary productivity and consumers up the food chain are also likely impacted. Furthermore, spongivorous fishes are, perhaps, found in higher abundance in these areas as they require sponges as a food source. Lastly, due to sponge’s ability to add to reef cementation, reefs where sponges are present in larger numbers may be more resilient to disturbances such as tropical cyclones. It is important to note that sponge size and abundance are the two factors which, together, determine the filtration intensity of sponges. Here we can directly associate abundance with ecological functional intensity since sponge size did not significantly differ across reefs and sites.

As the IPCC’s 2014 report predicts ocean temperatures to increase between 1-5 degrees Celsius, it is worth thinking about how ecologically and medically important taxa, such as sponges, may respond to these changes. From my filtration experiments, it is obvious that temperatures impact sponges physiologically. However, what is unclear is whether the patterns we see over a short time period, increased filtration efficiency with increased temperature, will continue over weeks, months, and even years. As ocean temperatures increase sponges may die as a result of too much physiological stress, adjust and revert back to “normal” filtration and physiological function, or simply sustain a higher filtration rate—thus changing the magnitude of their ecological impact as filter feeders. Further studies exploring filtration over longer time periods would help parse out some of these questions.

Sponges are not only an important reef contributor, they are also an invaluable medical resource since many sponges have bioactive compounds that contain anti-bacterial, anti-cancer, and anti-inflammatory properties (Murti and Agrawal 2010). For these reasons, it is critical to understand, first, where sponges are currently distributed and what this can then tell us about their habitat preferences/physiological tolerances. And, second, how they are likely to be impacted by changing climate conditions. My study laid down the foundation for future studies to address more specific questions about the habitat preferences and physiology of *Lamellodysidea*. Additionally, this study provided the knowledge necessary to track these sponge’s range over the coming years. Sponges are just one of the many groups of species which provide valuable ecosystem services and participate in important species interactions in coral reefs. Understanding the distribution and resilience of sponges to disturbances will be critical to predict how they will deal with changing conditions and thus function in future reef ecosystems.

## Acknowledgments

Thank you to all the professors—Brent Mishler, George Roderick, Stephanie Carlson, and Seth Finnegan—for the enormous amount of time and energy you put into this course. Thank you, Ilana Stein, Mo Tatlhego, and Phil Georgakakos for all your help with project design and R. Thank you to all my buddies, especially Madi Alcalay and Lucas Kampman for assisting with field work. And thank you class of 2019 for making this such a memorable experience.

## APPENDIX A

### Habitat Characterization

#### Fringing Reef

The first reef out from the shore until the shelf break or where coral becomes sparser again. Reef characteristics vary from very built-up to close, but interspersed coral. The shelf area was also included as part of the fringing reef zone. Transects were not conducted in the first 10m of the fringing reef such that the first GPS point was checked at 10m and it was determined whether a transect would be placed.

#### Mid-barrier

This area is described as the zone between the end of the fringing reef, often delineated as the north side of the channel, and the beginning of the barrier reef. A sandy bottom with interspersed coral is characteristic of this habitat. The range of coral density varies from very sparse with less than five coral heads along my transect to coral covering around 60% of the ground. This habitat is 350m or farther South from crest.

#### Barrier

Starting no sooner than 350m from the crest, the barrier region is characterized as having high coral and/or coral rubble ground cover with few to no sandy patches. This zone ends at the crest.

## APPENDIX B

**FIG. 1B.**
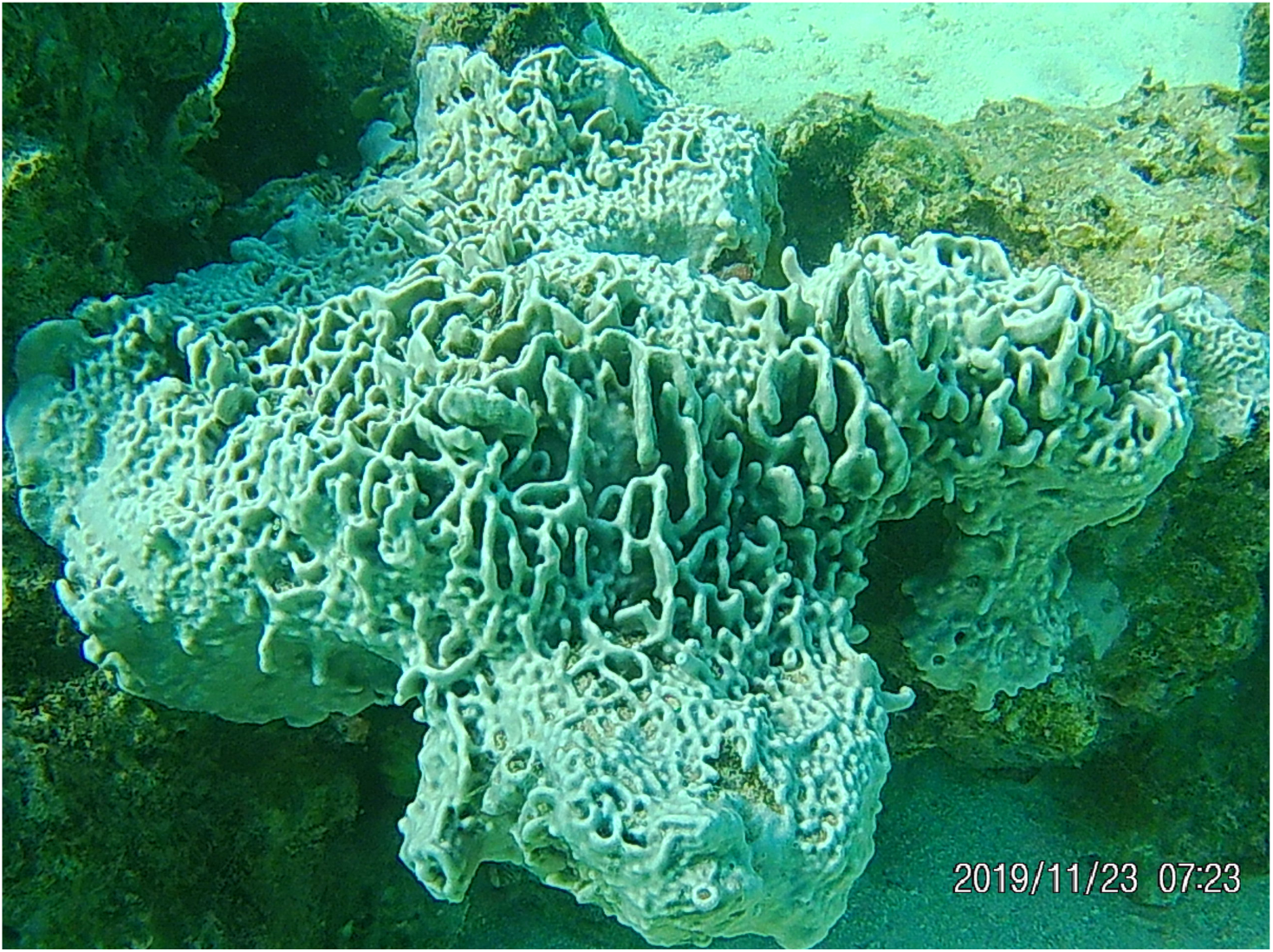
depicts the white/grey color variant of *Lamellodysidea sp..* The morphology here would be described as encrusting/tall because it displays both a thin encrusting form as well as tall projections.

**FIG. 2B.**
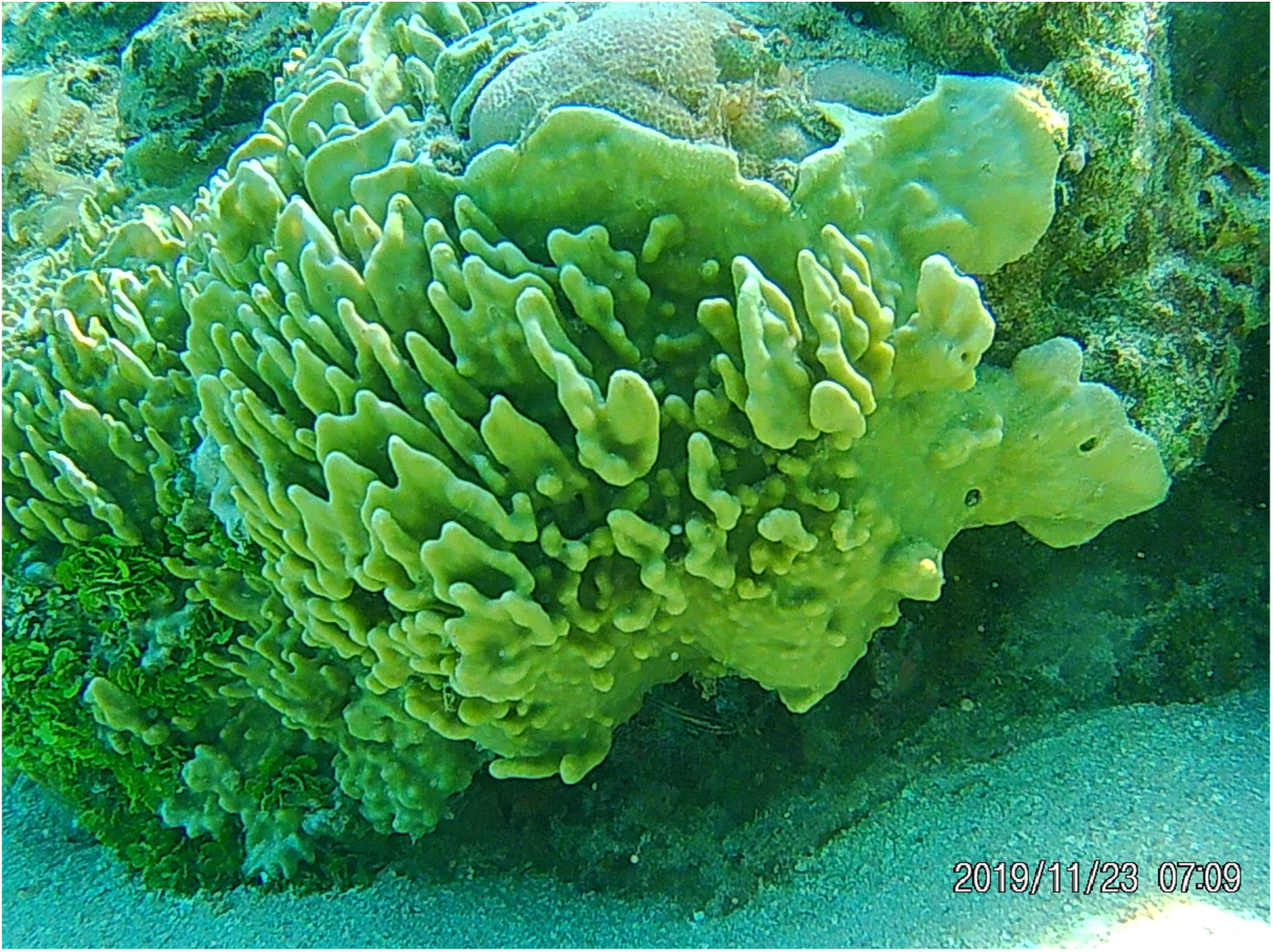
Shows the white/yellow variant of *Lamellodysidea sp.* This would also be considered encrusting/tall.

**FIG. 3B.**
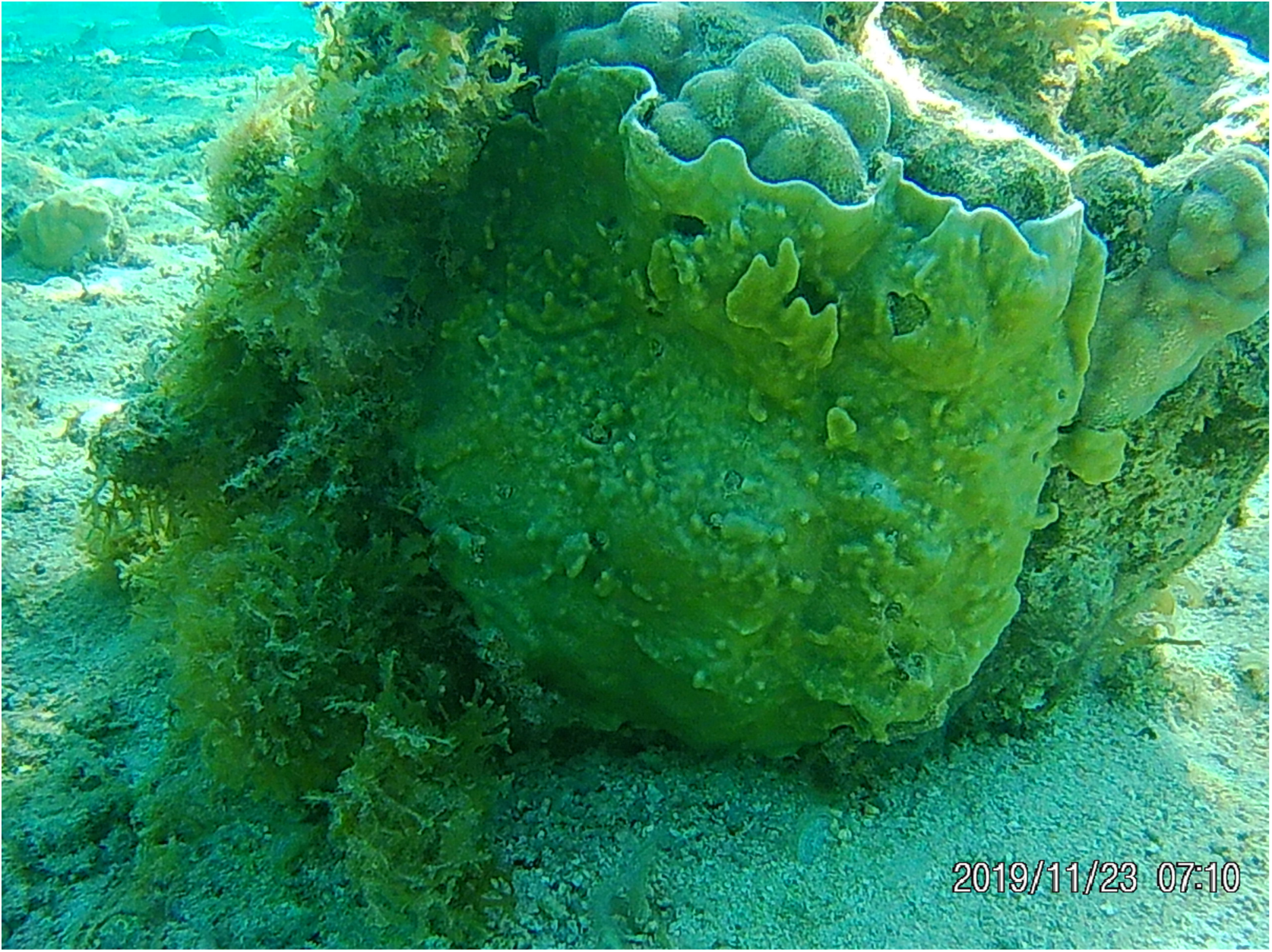
This sponge would be categorized as encrusting and white/yellow.

**Fig 4B.**
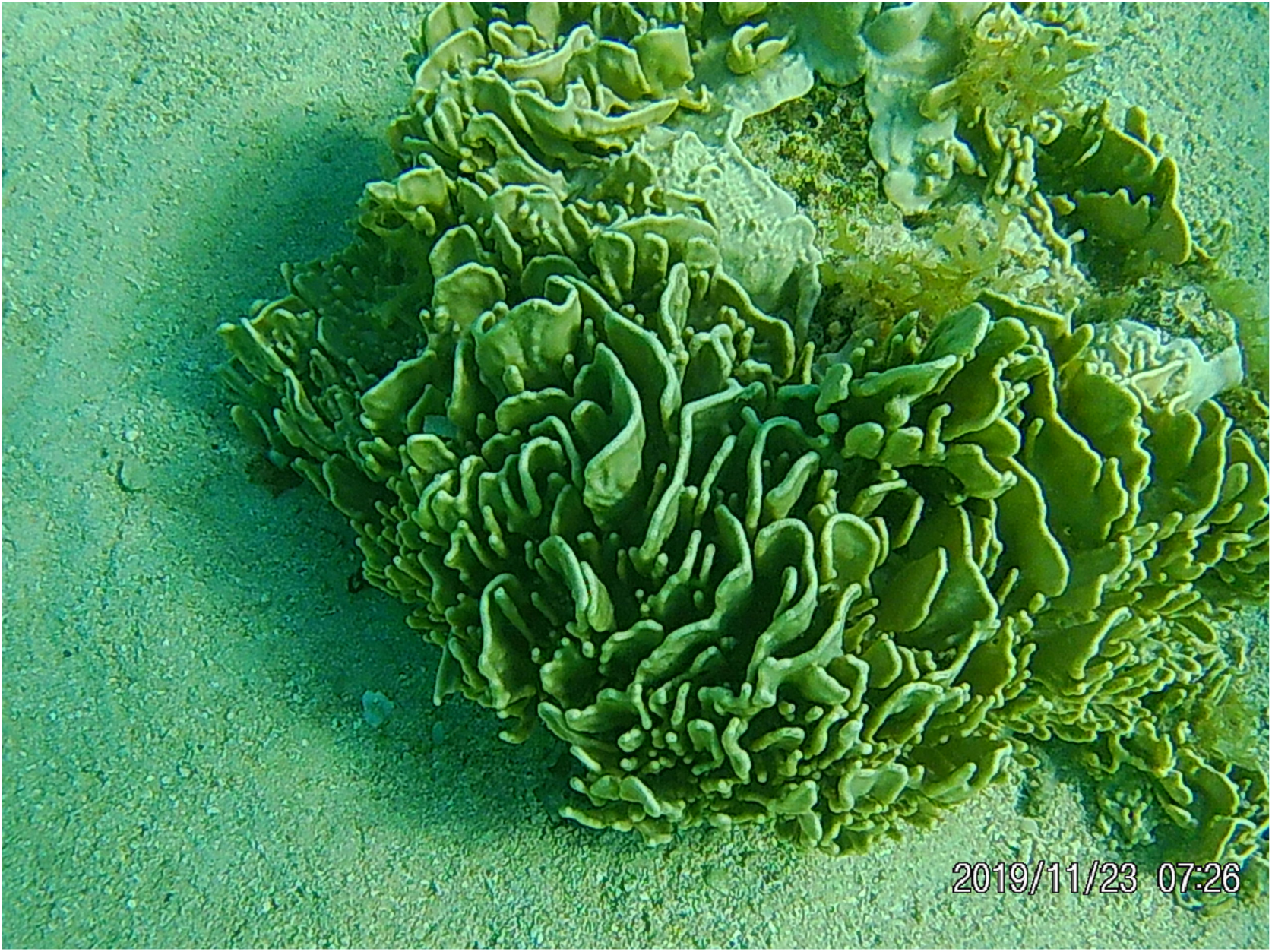
Shows a tall white/yellow sponge.

**Fig 5B.**
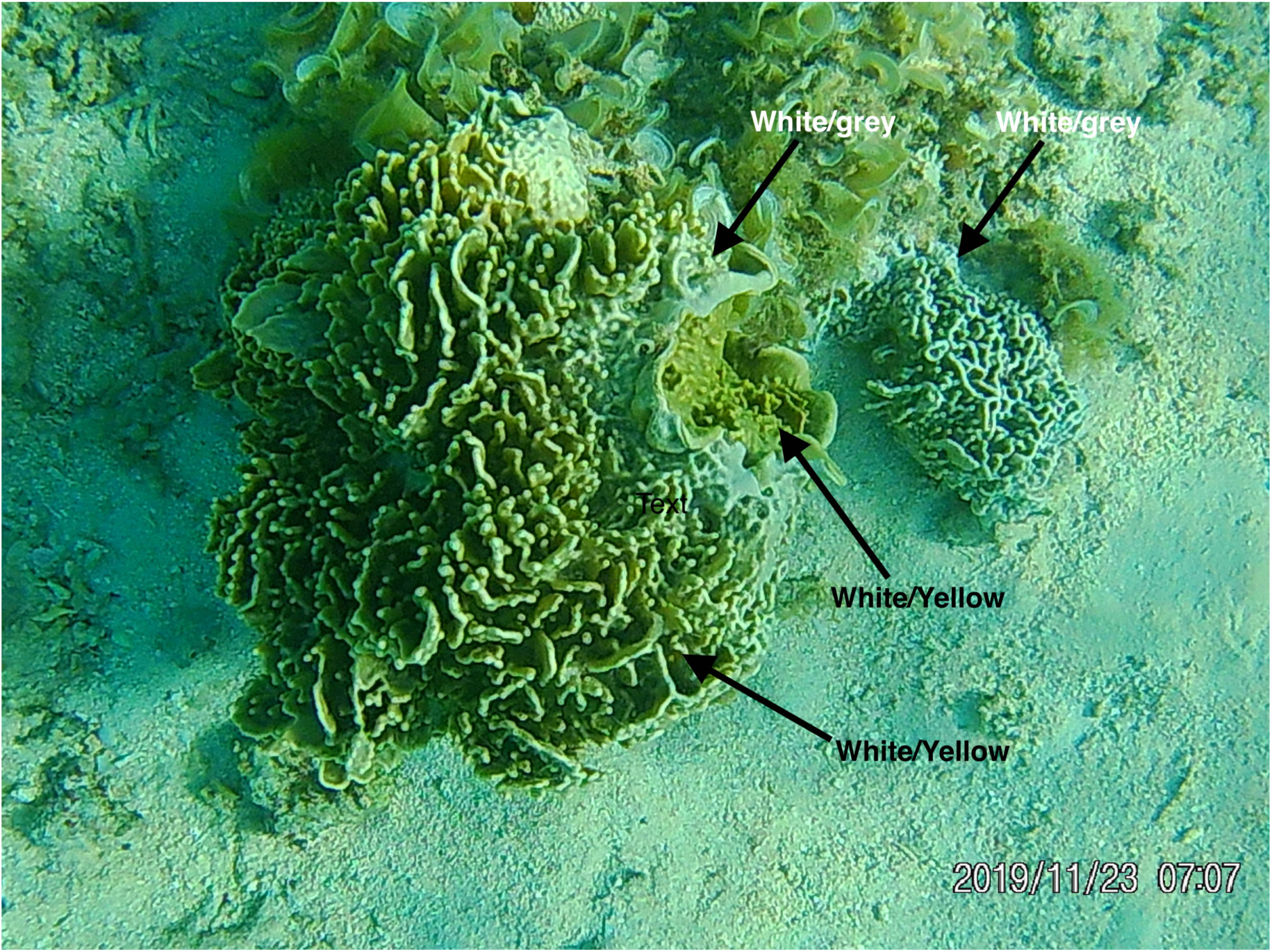
This shows how an individual sponge can contain sections of different color variants.

## APPENDIX C

| Site | Habitat | Transect | GPS Coordinates |
| --- | --- | --- | --- |
| 1 | Fringing | 1, 2, 3 | (1) 17°29'15.3"S, 149°49'34.1"W<br>(2) (2) 17°29'14.9"S, 149°49' 33.5" W<br>(3) 17°29'12.6"S, 149°49' 30.0"W |
| 1 | Mid-barrier | 1, 2, 3 | (1) 17°28'56.0"S,149°49'34.4"W<br>(2) 17°28'53.5"S, 149° 49' 35.6"W<br>(3) 17°28'52.7"S,149° 49' 36.1"W |
| 1 | Barrier | 1, 2, 3 | (1) 17°28'49.0"S,149°49' 37.9"W<br>(2) 17°28'47.7"S,149° 49' 38.1"W<br>(3) 17°28'47.1"S, 149° 49' 38.5"W |
| 2 | Fringing | 1, 2, 3, | (1) 17°29'6.7"S, 149°50'6.1"W<br>(2) 17°29'5.8"S, 149°50'6.1" W<br>(3) 17°29'5.6"S, 149°50'5.9"W |
| 2 | Mid-barrier | 1,2,3 | (1) 17°29'0.3"S, 149°50'5.4"W<br>(2) 17°28'55.8"S, 149°50'5.8"W<br>(3) 17°28'50.45"S, 149°50'5.0"W |
| 2 | Barrier | 1, 2, 3 | (1) 17°28'45.1"S, 149°50'6.5"W<br>(2) 17°28'41.0"S, 149°50'6.2"W<br>(3) 17°28'46.55"S, 149°50'5.8"W |
| 3 | Fringing | 1, 2, 3 | (1) 17°29'1.7"S, 149°50'31.3"W<br>(2) 17°29'0.20"S, 149°50'31.4"W<br>(3) 17°28'56.7"S, 149°50'30.9"W |
| 3 | Mid-barrier | 1, 2, 3 | (1) 17°28'54.6"S, 149°50'31.6"W<br>(2) 17°28'50.7"S,149°50'31.0"W<br>(3) 17°28'48.45"S,149°50'32.5"W |
| 3 | Barrier | 1, 2, 3 | (1) 17°28'43.6"S,149°50'33.8"W<br>(2) 17°28'41.5"S,149°50'33.6"W<br>(3) 17°28'39.7"S,149°50'33.1"W |
| 4 | Fringing | 1, 2, 3 | (1) 17°29' 24.8"S,149°50'58.4"W<br>(2) 17°29'24.5"S,149°50'58.6"W<br>(3) 17°29'23.6"S,149°50'59.3"W |
| 4 | Mid-barrier | 1, 2, 3 | (1) 17°29'17.2"S,149°51'6.1"W<br>(2) 17°29'12.9"S, 149°51'6.1"W<br>(3) 17°29'19.2"S,149°51'5.7"W |
| 4 | Barrier | 1, 2, 3 | (1) 17°29'10.6"S 149°51'11.9"W<br>(2) 17°29'07.8"S 149°51'14.8"W<br>(3) 17°29'05.4"S 149°51'16.4"W |
| Sponge collection site: |  |  | 17°28'52.1"S 149°49'59.5"W |

